# Estimating the seroprevalence of tuberculosis (*Mycobacterium bovis*) infection in a wild deer population in southwest England

**DOI:** 10.1101/2024.10.03.613747

**Authors:** Rachel Jinks, Alison Hollingdale, Rachelle Avigad, Juan Velarde, Chris Pugsley, Ricardo de la Rua-Domenech, Charlotte Pritchard, Tony Roberts, Julia Clark, Nick Robinson, Ruth Maynes, Susan Withenshaw, Graham Smith

**Author notes:** **Corresponding author:** Rachel Jinks.

## Abstract

Bovine tuberculosis (bTB) is a major disease of cattle that is subject to an eradication strategy in England. To inform control policies and manage the epidemic, all potential sources of infection for cattle must be identified and understood. The causative agent of bTB, *Mycobacterium bovis*, has a wide host range including several deer species. While transmission between cattle and deer has been implicated in some localised endemic regions, the role of deer in the epidemiology of bTB in England is poorly understood. This paper presents the results of a serological survey to estimate the prevalence of *M. bovis* in a large wild deer population in the High bTB Risk Area of southwest England.

Blood samples were collected post-mortem over a 12-month period from wild deer during annual deer management controls in the Exmoor area and tested for *M. bovis* serum antibodies. Overall, 432 samples were collected and 69 (16.0%) were seropositive. The true seroprevalence in our sample was estimated to be 29.2% (95% CrI 21.1-38.6%), using Bayesian Latent Class Analysis to account for imperfect diagnostic test accuracy. Prevalence did not appear to differ between sexes nor between species, although the sample was predominantly red deer. The lowest prevalence was observed in animals aged under 1-year.

Whilst these results provide valuable insights into *M. bovis* seroprevalence in this wild deer population, they should be interpreted alongside other relevant information such as ecology, species-specific epidemiology and disease pathology, deer density and cattle management to inform potential transmission risks between cattle and wild deer.

## INTRODUCTION

Bovine tuberculosis (bTB) is a chronic infectious disease of cattle caused by the bacterium *Mycobacterium bovis* (*M. bovis*), which can also infect and give rise to TB in a wild range of other hosts including humans, badgers, deer, and other mammalian species. The epidemiology of bTB is complex, particularly in areas where both cattle and wildlife hosts reside.

Bovine TB has been described as the most pressing animal health problem in England (Defra, 2014). This is due to the economic, social, and animal welfare impacts that the disease and the measures to control it have on affected animals, herds, the wider livestock industry, wildlife, and international trade.

Combined annual costs of the bTB eradication strategy to the farming industry and the UK government amount to approximately £150 million a year (Defra, 2018). This includes cattle and wildlife surveillance and control measures, compensation payments to keepers of animals removed for bTB control reasons, and government-funded research and development. The disease has been subject to a statutory eradication program in Great Britain (GB) since 1950. In 2014, the UK government published a strategy for achieving Officially Bovine Tuberculosis Free status for England by 2038 (Defra, 2014). This strategy consists of several interventions to be taken towards eradication, focusing on more stringent surveillance and control measures in cattle, supplemented by badger controls in areas of endemic infection in England. An independent review of the strategy completed in 2018 made a series of recommendations to accelerate disease eradication, including enhancing knowledge of *M. bovis* infection prevalence in other susceptible wildlife species (Godfray et al., 2018). In August 2024, the new UK government announced a refreshed TB eradication strategy, including a national wildlife surveillance programme.

For bTB eradication purposes, England is divided into three ‘risk’ areas depending on the prevalence and epidemiology of the disease - the High Risk area (HRA), the Low Risk area (LRA) and the Edge Area (EA), with different cattle and wildlife disease control measures in each (Animal and Plant Health Agency, 2022). The HRA, which includes the southwest of England, has historically had the highest incidence rate and prevalence of bTB in cattle herds in England (Animal and Plant Health Agency, 2022). From 2011 to 2018, the incidence rate for cattle herds in the HRA fluctuated between 18 and 20 new incidents per 100 herd-years at risk (HYR) but has been decreasing since. In 2023, the herd incidence rate in the HRA was the lowest recorded since 2006, at 13.2 incidents per 100 HYR (Animal and Plant Health Agency, 2024)

The bTB epidemic in England has been in decline over the last six years (Animal and Plant Health Agency, 2022), but all potential sources of infection and risk pathways for cattle herds must be identified and understood to meet the eradication timeframe. In GB, the European badger *(Meles meles)* has been identified as the main wildlife reservoir (Godfray et al., 2013, Krebs et al., 1997). Since 2013, badger culling licences have been granted in the HRA of England as part of the government’s bTB eradication strategy (Defra, 2014), alongside further cattle control measures. The role of badgers in bTB transmission has been more regularly studied than other wildlife hosts (Justus et al., 2024). However, previous studies have also identified other British wild mammals as being infected with *M. bovis*, including several species of deer in the southwest region of England (Collard, 2023, Delahay et al., 2007). Delahay et al. (2007) performed a semi-quantitative risk assessment of alternative wildlife hosts based on their *M. bovis* infection prevalence, typical pathology, ecology, and density, and concluded that the risk to cattle from deer species was lower than that posed by badgers, although potentially substantial, particularly for fallow (*Dama dama*) and red (*Cervus elaphus*) deer. As there are wide variations in deer density and distribution across the southwest of England, the estimated risk to cattle herds from infected deer populations is likely to be localised, with a very high level of uncertainty (Delahay et al., 2007).

Considering the previous semi-quantitative assessment of the risk posed to local cattle herds by wild deer species, we conducted this study to increase knowledge of *M. bovis* in wild deer populations in an area of endemic bTB herd incidence in England. Our aim was to use opportunistically collected data from annual deer management culls to provide an estimate of *M. bovis* prevalence in a large wild deer population in the HRA area of England, where sporadic cases of TB in wild deer carcases had been documented for several years.

## MATERIALS AND METHODS

### Study area

The study area, known as Exmoor, straddles northern Devon and Somerset, two counties in southwest England. Much of Exmoor was designated as a National Park in 1954, a large area of land protected by national legislation (National Parks UK, 2024). Exmoor National Park covers an area of approximately 267 square miles (692km^2^), two thirds of which are in Somerset and one third in Devon (Figure 1). Land type in Exmoor consists of high moorland, 34 miles of steep coastline, ancient woodland, and agricultural land. The Agricultural Land Classification (ALC) system grades most of this area as Grade 4 (poor) or Grade 5 (very poor) (Natural England, 2010). The predominant agricultural use of land is grazing for sheep and cattle (Natural England, 2012).

**Figure 1.**
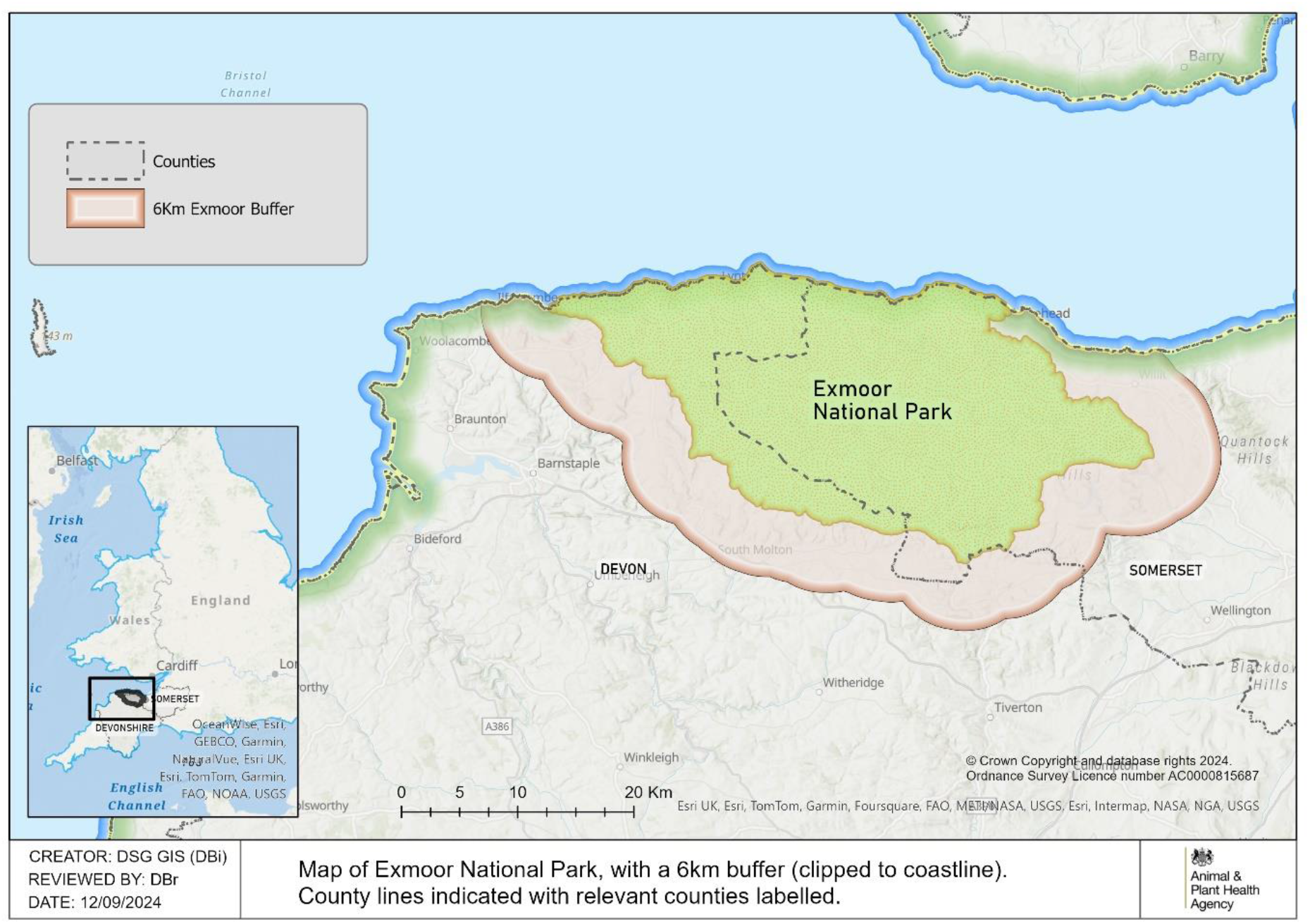
Map of Exmoor National Park with a 6km buffer delineating the extent of the study area for wild deer sampling purposes

Exmoor also maintains a large population of wild deer, which are free to roam over the whole area and beyond. Since 1994, there has been an annual deer count census on Exmoor, managed by a local stakeholder group, which has estimated the red deer population to be approximately 3700 in recent years (unpublished data from the 2022 and 2023 deer census). There are also much smaller populations of other deer species inhabiting the area, including fallow, roe *(Capreolus capreolus*) and muntjac *(Muntiacus reevesi*). All samples submitted for this study were collected from carcases of wild deer shot within an area spanning Exmoor National Park and a 6km buffer around the park boundary (Figure 1).

### Sample collection

A single blood sample (5-10ml) was collected immediately post-mortem from wild deer shot during annual wild deer management controls in the study area (Figure 1). Samples were collected by local deer managers (local individuals or organisations) over a one-year period and provided to APHA. Samples were collected between 1^st^ August 2022 and 31^st^ July 2023, observing the wild deer species-specific open and closed seasons in England (The British Deer Society, 2024).

Samples were transported from the field to a local central collection point as soon as possible after collection and refrigerated. Samples were then couriered to the nearest APHA Regional Laboratory within seven days of collection. Once received at the APHA Regional Laboratory, the blood samples were spun at 672g for ten minutes to obtain serum. The serum was frozen and stored at -80°C, before thawing and testing as per internal Standard Operating Procedures (SOP).

Blood sample tubes were individually identified at collection and submitted with basic demographic data on the location (as a grid reference or northing / easting), sex, species, and estimated age of the shot animal. The visual condition and any obvious external injuries of the animal were recorded as a free text description for some of the samples.

### Laboratory testing

The IDEXX *M. bovis* ELISA test is a commercial kit to detect cattle TB antibodies to two immunodominant mycobacterial antigens (MPB70 & MPB83). The assay has been modified by APHA to detect cervine antibodies (Barton et al., 2023). The serum samples were tested following the protocol described in Barton et al. (2023) and internal APHA Standard Operating Procedures. Up to 90 samples were tested per ELISA plate, together with plate positive and negative quality controls. The deer IDEXX antibody test has a specificity of 98.8% (95% CI: 97.2-99.6%) and a moderate sensitivity of 55.2% (45.2-65.0%) when no prior skin test is performed (non-anamnestic samples, as was the case with this study), using the cut off >0.502 for a positive result (Barton et al., 2023).

### Statistical methods

The number of positive samples on the IDEXX antibody test gives a crude estimate of the apparent seroprevalence in the target population. However, misclassification is possible due to the imperfect diagnostic accuracy of the test used and so this estimate should be adjusted accordingly. The IDEXX test for deer has a sensitivity of 55.2% on non-anamnestic samples (Barton et al., 2023), so the true seroprevalence is likely to be higher than the apparent seroprevalence. We used Bayesian Latent Class analysis (BLCA) to derive an estimate of the true seroprevalence of *M. bovis* (the latent class) based on the variables for which estimates do exist (apparent prevalence, and sensitivity and specificity of the IDEXX test) (Branscum et al., 2005). While the point estimate for true prevalence should be broadly the same whether BLCA or a more traditional method of adjustment (such as the Rogan-Gladen estimator) is used, the inclusion of prior distributions rather than single values for sensitivity and specificity in BLCA results in improved confidence interval coverage compared to traditional methods (Flor et al., 2020).

The analyses were implemented in JAGS via the runjags package (Denwood, 2016) in R version 4.4.0 (R Core Team, 2024). Chains were run for 10,000 iterations, after a burn in of 6,000 (discarded). Each parameter history was checked graphically to ensure convergence, and three different starting values were used for chains to ensure the results were robust to different starting values of the parameters. Informative prior distributions were used. For sensitivity and specificity of the IDEXX test the priors were based on the only estimates of test performance currently available (Barton et al., 2023) with the median sensitivity slightly reduced to account for the fact that in Barton et al the gold standard (mycobacterial culture) was assumed to be a perfect test. The priors for prevalence of *M. bovis* were based on results published recently also studying the Exmoor wild deer population(Collard, 2023).

## RESULTS

### Demographics of the surveyed deer

In total, blood samples were received from 432 wild deer, including males and females of varying ages, from red, roe, fallow and muntjac species. The sex, age and species distribution of the samples are given in Table 1.

**Table 1.** Distribution of sex, age and species for deer included in the study sample.

|  | N animals | % of sample |
| --- | --- | --- |
| **Total** | 432 | 100.0% |
| **Sex** |  |  |
| Female | 225 | 52.1% |
| Male | 206 | 47.7% |
| Not recorded | 1 | 0.2% |
| **Age category \*** |  |  |
| Under 1 | 37 | 8.6% |
| 1-2 years | 37 | 8.6% |
| 2-3 years | 84 | 19.4% |
| 3-4 years | 90 | 20.8% |
| 4-6 years | 100 | 23.1% |
| 6-8 years | 32 | 7.4% |
| 8-10 years | 20 | 4.6% |
| 10+ years | 15 | 3.5% |
| Not recorded | 17 | 3.9% |
| **Species** |  |  |
| Red | 347 | 80.3% |
| Fallow | 63 | 14.6% |
| Roe | 19 | 4.4% |
| Muntjac | 1 | 0.2% |
| Not recorded | 2 | 0.5% |

\*Age: category named 'x-y years' includes animals estimated to be at least x years old, but not yet y years old

Samples were received from across the Exmoor area, including up to 6km from the outer National Park boundary, as shown in Figure 1. No samples were received from some of the westernmost parts of the study area, due to limited stakeholder engagement in these areas.

The data regarding deer body condition was not of sufficient quality to be formally included in the analysis. However, it showed that the wild deer contributing samples to the study included animals with varying external body conditions, ranging from externally ‘healthy’ looking, to injured animals (for example broken leg), to visually unhealthy animals (for example very poor body condition). Whilst it is difficult to establish if a truly representative population sample is tested, due to the nature of the wild and free roaming subjects and the opportunistic nature of the sampling, the data collected suggests the study did capture a broad representation of the population. The identity of the local deer manager responsible for each sample submitted was recorded. In total, 25 different individuals or organisations contributed to this study. Ten individuals or organisations contributed 10 or more study samples and were responsible for submitting 375 samples (86.8% of the total). Again, the range of contributors to the study was broad and each were managing the deer population for varying reasons, including for food production, to alleviate animal suffering and to manage the deer numbers.

### *M. bovis* seroprevalence in surveyed deer samples

Table 2 presents the number and percentage of IDEXX positive samples, and the estimated true seroprevalence of *M. bovis* calculated using BLCA, by sex, age group and deer species. Overall, 69 (16%) out of 432 samples were positive on the IDEXX test and the true seroprevalence was estimated to be 29.2% (95% credible interval 21.1 - 38.6%). Males and females had a very similar seroprevalence of 29.7% (20.0 - 40.3%) and 29.6% (19.7 - 40.6%). None of the blood samples from wild deer under 1 year old were positive on the IDEXX test, leading to an estimated seroprevalence of 18.0% (7.0 - 38.6%) once the low sensitivity of the test was accounted for. The age group with the highest seroprevalence was the 2-3 year-olds at 35.0% (22.3 - 49.4%), followed by the 10+ year olds 33.1% (16.0 - 51.3%). The other age groups ranged between 29.0% and 31.3%. Most samples (347/432) came from red deer, which had an estimated seroprevalence of 28.7% (20.1 - 38.1%). Fallow deer had a similar level. A slightly higher level was observed in roe deer at 33.8% (17.7 - 52.0%); however, this was a small sample (n=19).

**Table 2.** Apparent and estimated true *M. bovis* seroprevalence overall and for sex, age, and species subgroups.

|  | **N IDEXX positive /**<br>**N animals sampled** | **% animals IDEXX positive**<br>**(apparent seroprevalence)** | **True seroprevalence estimate**<br>**using BLCA (median, 95% CrI)** |
| --- | --- | --- | --- |
| **Total** | 69 / 432 | 16.0% | 29.2% (21.1 - 38.6%) |
| **Sex** |  |  |  |
| Female | 36 / 225 | 16.0% | 29.6% (20.0 - 40.3%) |
| Male | 33 / 206 | 16.0% | 29.7% (19.7 - 40.6%) |
| Not recorded | 0 / 1 | 0.0% |  |
| **Age category** |  |  |  |
| Under 1 | 0 / 37 | 0.0% | 18.0% (7.0 - 32.0%) |
| 1-2 years | 5 / 37 | 13.5% | 29.0% (14.9 - 45.3%) |
| 2-3 years | 17 / 84 | 20.2% | 35.0% (22.3 - 49.4%) |
| 3-4 years | 14 / 90 | 15.6% | 29.7% (17.6 - 43.2%) |
| 4-6 years | 17 / 100 | 17.0% | 31.3% (19.4 - 44.4%) |
| 6-8 years | 5 / 32 | 15.6% | 30.8% (16.2 - 47.4%) |
| 8-10 years | 3 / 20 | 15.0% | 30.8% (15.0 - 48.0%) |
| 10+ years | 3 / 15 | 20.0% | 33.1% (16.0 - 51.3%) |
| Not recorded | 5 / 17 |  |  |
| **Species** |  |  |  |
| Red | 54 / 347 | 15.6% | 28.7% (20.1 - 38.1%) |
| Fallow | 10 / 63 | 15.9% | 30.3% (16.9 - 44.6%) |
| Roe | 4 / 19 | 21.1% | 33.8% (17.7 - 52.0%) |
| Muntjac | 0 / 1 | 0.0% | 31.6% (14.1 - 51.9%) |
| Not recorded | 1 / 2 | 50.0% |  |

To look more closely at the percentage of *M. bovis* seropositive samples across age ranges, the true seroprevalence was also estimated for each age category separately for males and females (Table 3). These estimates are illustrated in Figure 2, which shows there was no clear difference in seroprevalence between the sexes. However, the numbers in each subgroup were too small to draw firm conclusions, especially since the estimation of age in the adult animals was not precise.

**Table 3.** Apparent and estimated true *M. bovis* seroprevalence for age subgroups within Males and Females.

|  | **N IDEXX positive /**<br>**N animals sampled** | **% samples IDEXX positive**<br>**(apparent seroprevalence)** | **True seroprevalence estimate**<br>**using BLCA (median, 95% CrI)** |
| --- | --- | --- | --- |
| **Females** |  |  |  |
| Under 1 | 0 / 22 | 0.0% | 22.1% (8.4 - 38.1%) |
| 1-2 years | 2 / 12 | 16.7% | 31.9% (15.2 - 50.5%) |
| 2-3 years | 9 / 38 | 23.7% | 36.8% (21.6 - 53.2%) |
| 3-4 years | 9 / 48 | 18.8% | 33.0% (19.0 - 48.7%) |
| 4-6 years | 10 / 61 | 16.4% | 30.9% (17.9 - 45.8%) |
| 6-8 years | 2 / 22 | 9.1% | 27.2% (12.2 - 44.2%) |
| 8-10 years | 1 / 8 | 12.5% | 31.0% (14.3 - 50.2%) |
| 10+ years | 1 / 2 | 50.0% | 34.4% (16.2 - 54.1%) |
| Not recorded | 2 / 12 |  |  |
| **Males** |  |  |  |
| Under 1 | 0 / 14 | 0.0% | 25.1% (10.1 - 42.7%) |
| 1-2 years | 3 / 25 | 12.0% | 28.8% (14.2 - 45.7%) |
| 2-3 years | 8 / 46 | 17.4% | 32.0% (17.4 - 47.4%) |
| 3-4 years | 5 / 42 | 11.9% | 27.2% (13.6 - 42.5%) |
| 4-6 years | 7 / 39 | 17.9% | 32.4% (17.8 - 48.8%) |
| 6-8 years | 3 / 10 | 30.0% | 35.9% (18.4 - 54.8%) |
| 8-10 years | 2 / 12 | 16.7% | 31.9% (14.9 - 50.1%) |
| 10+ years | 2 / 13 | 15.4% | 31.5% (15.2 - 50.4%) |
| Not recorded | 3 / 5 |  |  |

**Figure 2.**
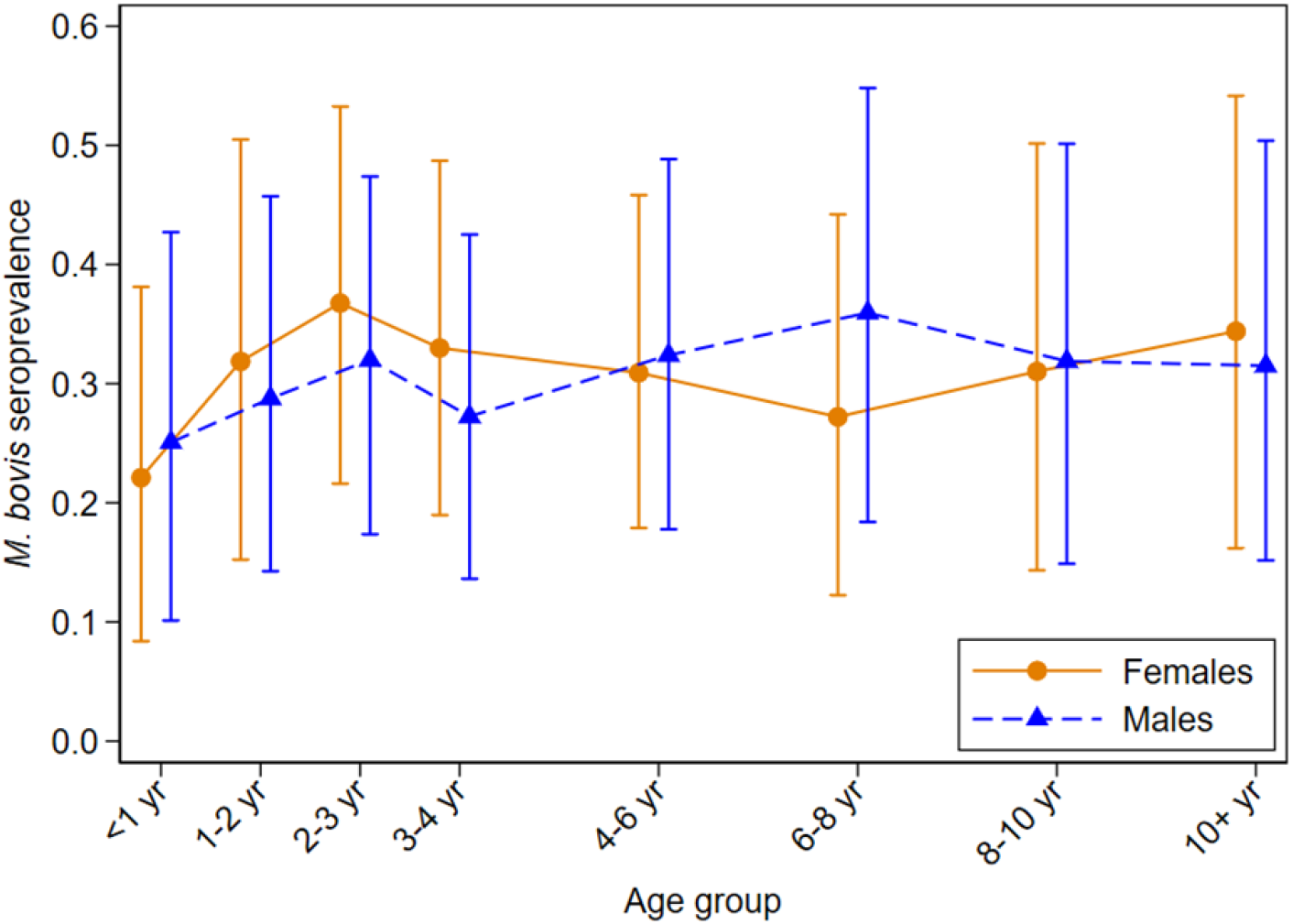
Estimated true *M. bovis* seroprevalence by age for female and male deer, with 95% credible intervals.

## DISCUSSION

This study is, to our knowledge, the largest *M. bovis* seroprevalence survey conducted in the Exmoor wild deer population to date. Blood samples were collected across the study area over a 12-month period, plus the large number of different contributors helped make the study sample a broad representation of this wild deer population. However, as the samples were obtained opportunistically and voluntarily, a degree of sampling bias in the study results cannot be excluded. The results provide an insight into the current seroprevalence of *M. bovis* in wild deer in this area of southwest England. The estimated overall seroprevalence in our sample was 29.2% (21.1 - 38.6%), using Bayesian latent class analysis to adjust for the imperfect diagnostic accuracy of the IDEXX test. The seroprevalence did not appear to differ between males and females in this study and there was little evidence of differences between deer species, although the study sample consisted predominantly of red deer. There was no clear relationship between age and seroprevalence beyond the apparently much lower level observed for the youngest animals.

Our findings are similar to those reported in a recent smaller (n=106) field survey of wild red deer in Exmoor (Collard, 2023). In that study, the cervid DPP Vet TB kit was used, and the prevalence estimates were not adjusted for potential false negative and false positive test results, although sensitivity of the test was reported to be much higher than the IDEXX test at up to 91%. Overall prevalence was estimated to be 28.3% and little difference was seen between males and females (Collard, 2023). A much higher seroprevalence (42.9%) was observed in the youngest age category (‘yearlings’) in Collard’s sample, compared with 18.0% adjusted prevalence in our study in the under 1 year-old group and 29.0% in the 1-2 year group. However, it is not clear whether the age categories used by Collard are equivalent to those used in our analysis. Studies of red deer in other locations have also found low prevalence of *M. bovis* infection in the youngest animals: a study in New Zealand found that only 1 / 18 animals under 1 year old in their sample showed histological evidence of TB (Lugton et al., 1998), while a large study in Northern Spain of 2759 ‘hunter-harvested’ red deer estimated the prevalence to be ∼3-4% in calves (under 1 year) and ∼8% for yearlings (in their second season) (Vicente et al., 2013). However, the assessments of prevalence in these two studies were based on culture or visible lesion methodologies rather than serology, and so may not be directly comparable to our study.

The accuracy of our true seroprevalence estimates was hampered by the limited evidence available on the sensitivity and specificity of the IDEXX test in deer. Estimates of the diagnostic accuracy of this test are currently based on one publication (Barton et al., 2023), and if its sensitivity is less than the estimated 55%, then the true prevalence of *M. bovis* infection in Exmoor wild deer could be higher than reported in this study. We mitigated this by incorporating a slightly lower median in our prior distribution for sensitivity in the BLCA, so our results should be robust to the test sensitivity being up to 3% lower. As part of the model development process, we also considered a prior distribution with median IDEXX sensitivity as low as 50%, and with wider confidence intervals, and found that even with this poorer performance, the upper limit of the 95% credible interval for overall prevalence did not exceed 50%.

The overall seroprevalence of 29.2% in this study would lie at the upper range of worldwide pooled prevalence estimates of TB in global red deer populations (Reis et al., 2021). Given the location of our study area in the HRA of England, regular notifications of tuberculous wild deer carcases in Exmoor through statutory passive surveillance, and the endemic nature of bTB in this area, the overall prevalence detected in the local wild deer population was not a wholly unexpected finding. According to the meta-analysis conducted by Reis et al. (2021) which considered research using both serological and tissue analysis, the countries with the highest estimated pooled prevalence of TB in red deer were Austria (31.6%) and Portugal (27.8%). Somecountries showed a moderate prevalence in red deer (France 6.4%; Spain 12.1%), whilst others showed lower levels (Switzerland 0.2%; UK 1.0%). An analysis of mean TB prevalence in European wildlife species estimated a prevalence of 20% in fallow deer (Justus et al., 2024). Delahay et al. (2007) estimated the TB prevalence in UK wild deer to be 1.0% in roe and red deer, 4.4% in fallow deer and 5.2% in muntjac, although this study also stated that their figures may have been underestimated due to the sampling method. Delahay et al. (2007) also concluded that deer should be considered as a potential source of infection for cattle herds in localised areas of the UK. Further work to understand prevalence rates in other wild deer populations across the HRA would be beneficial.

However, caution should be used in interpreting data on the prevalence of infection in wildlife populations in isolation. Prevalence data should be interpreted with other relevant information such as ecology, species-specific epidemiology and disease pathology, deer density and local cattle management to inform potential transmission risks to cattle herds. Our results highlight the importance of studying other wildlife species in bTB endemic areas, where they co-exist with cattle and other susceptible species that may be involved in the epidemiology of bTB. Multi-species surveillance techniques would be beneficial to further understand the complex epidemiology of *M. bovis* infections in multi-host ecosystems (Justus et al., 2024), as we work towards bTB eradication in England.

Our study was part of a wider initiative in the Exmoor area, to encourage the reporting of suspect lesions of TB in wild deer carcasses to APHA, which is a statutory requirement. A further aim of this initiative is to carry out Whole Genome Sequence (WGS) analysis of all isolates of *M. bovis*, from approximately 100 culture-positive wild deer tissue samples, by funding the collection, sampling, and appropriate disposal of the suspected tuberculous carcases (with funding provided by the Department for the Environment, Food and Rural Affairs, Defra). It is envisaged that the WGS results produced will be analysed to build time-stamped phylogenies of *M. bovis* isolates from local cattle, deer, and badgers, and apply models that leverage genetic information to estimate evolutionary dynamics, such as between-species transmission rates and direction of transmission between the main host species. Historical wild deer samples from the area will also be included in the analysis, extending the temporal window, and strengthening the temporal signal of the dataset. Collection of these data is still ongoing. This further investigation will aid our understanding of disease transmission between different species, supporting the bTB eradication programme both locally and nationally.

This study was delivered through collaborative working between the local community, stakeholders, and Government. By working together, this study has increased our knowledge regarding the potential role of wild deer in the epidemiology of bTB in the southwest of England. These collaborations should be valued by all interested parties as we work towards eradicating bTB in England, resulting in wildlife populations and a cattle industry which are healthy and sustainable, benefiting both the economy and wider society.

## ETHICS APPROVAL STATEMENT

No animals were specifically culled for this study, as the samples were collected postmortem. Thus, no specific ethical approval was required.

## ACKNOWLEDGEMENTS

The authors would like to thank the local deer managers for contributing the samples for this study, and the local stakeholder groups involved for providing administrative and logistical assistance. Funding was provided by the Department for Environment, Food and Rural Affairs (Defra) and the Scottish and Welsh Governments through project SB4510 for testing of blood samples. Defra funded the sample transportation costs.

## Notes

### Competing Interest Statement

The authors have declared no competing interest.

## REFERENCES

Animal and Plant Health Agency 2022. Bovine tuberculosis in England in 2022. Animal and Plant Health Agency.

Animal and Plant Health Agency 2024. Tuberculosis (TB) in cattle in Great Britain. Detailed statistics on Tuberculosis (TB) in cattle in Great Britain with regional breakdowns: Headline statistical dataset. https://www.gov.uk/government/statistical-data-sets/tuberculosis-tb-in-cattle-in-great-britain.

Barton, P., Robinson, N., Middleton, S., O’Brien, A., Clarke, J., Dominguez, M., Gillgan, S., Selmes, J. & Rhodes, S. 2023. Evaluation of Antibody Tests for Mycobacterium bovis Infection in Pigs and Deer. Vet. Sci., 10.

Branscum, A. J., Gardner, I. A. & Johnson, W. O. 2005. Estimation of diagnostic-test sensitivity and specificity through Bayesian modeling. Prev. Vet. Med., 68, 145–163.

Collard, K. J. 2023. A study of the incidence of bovine tuberculosis in the wild red deer herd of Exmoor. Eur. J. Wildl. Res., 69, 14.

Defra 2014. The Strategy for achieving Officially Bovine Tuberculosis Free status for England. Department for Environment, Food and Rural Affairs.

Defra 2018. Next steps for the strategy for achieving bovine tuberculosis free status for England. The government’s response to the strategy review, 2018. Department for Environment, Food and Rural Affairs.

Delahay, R. J., Smith, G. C., Barlow, A. M., Walker, N., Harris, A., Clifton-Hadley, R. S. & Cheeseman, C. L. 2007. Bovine tuberculosis infection in wild mammals in the South-West region of England: A survey of prevalence and a semi-quantitative assessment of the relative risks to cattle. Vet. J., 173, 287–301.

Denwood, M. J. 2016. runjags: An R Package Providing Interface Utilities, Model Templates, Parallel Computing Methods and Additional Distributions for MCMC Models in JAGS. J. Stat. Softw., 71, 1–25.

Flor, M., Weiß, M., Selhorst, T., Müller-Graf, C. & Greiner, M. 2020. Comparison of Bayesian and frequentist methods for prevalence estimation under misclassification. BMC Public Health, 20, 1135.

Godfray, H. C. J., Donnelly, C., Hewinson, G., Winter, M. & Wood, J. 2018. Bovine TB strategy review. Department for Environment Food and Rural Affairs.

Godfray, H. C. J., Donnelly, C. A., Kao, R. R., Macdonald, D. W., McDonald, R. A., Petrokofsky, G., Wood, J. L. N., Woodroffe, R., Young, D. B. & McLean, A. R. 2013. A restatement of the natural science evidence base relevant to the control of bovine tuberculosis in Great Britain. Proceedings of the Royal Society B: Biological Sciences, 280.

Justus, W., Valle, S., Barton, O., Gresham, A. & Shannon, G. 2024. A review of bovine tuberculosis transmission risk in European wildlife communities. Mammal Rev., 54, 325–340.

Krebs, J. R., Anderson, R. A., Clutton-Brock, T., Morrison, I., Young, D. & Donnelly, C. 1997. Bovine Tuberculosis in Cattle and Badgers. Report to the Rt Hon Dr Jack Cunningham MP.

Lugton, I., Wilson, P., Morris, R. & Nugent, G. 1998. Epidemiology and pathogenesis of Mycobacterium bovis infection of red deer (Cervus elaphus) in New Zealand. N. Z. Vet. J., 46, 147–156.

National Parks UK. 2024. What is a National Park? [Online]. Available: https://www.nationalparks.uk/what-is-a-national-park/ [Accessed September 2024].

Natural England. 2010. Regional Agricultural Land Classification Maps [Online]. Available: https://publications.naturalengland.org.uk/category/5954148537204736 [Accessed September 2024].

Natural England. 2012. National Character Area Profile: 145 Exmoor (NE342) [Online]. Available: https://publications.naturalengland.org.uk/publication/2303045 [Accessed September 2024].

R Core Team 2024. R: A Language and Environment for Statistical Computing.

Reis, A. C., Ramos, B., Pereira, A. C. & Cunha, M. V. 2021. The hard numbers of tuberculosis epidemiology in wildlife: A meta-regression and systematic review. Transbound. Emerg. Dis., 68, 3257–3276.

The British Deer Society. 2024. Deer Open Season [Online]. Available: https://bds.org.uk/information-advice/resources/deer-open-season/ [Accessed September 2024].

Vicente, J., Barasona, J. A., Acevedo, P., Ruiz-Fons, J. F., Boadella, M., Diez-Delgado, I., Beltran-Beck, B., González-Barrio, D., Queirós, J., Montoro, V., de la Fuente, J. & Gortazar, C. 2013. Temporal trend of tuberculosis in wild ungulates from Mediterranean Spain. Transbound. Emerg. Dis., 60, 92–103.

